# Long-read, whole-genome sequencing and chemotherapy response of two patient-derived organoids from a TP53- and KRAS-mutant ovarian carcinoma

**DOI:** 10.64898/2026.07.06.736185

**Authors:** Jae Rim Wendt, Kristin M. Adams, Ryan Moreno, Md Shahadat Hossan, Austin Stram, Ethan S. Lin, Lauren Kersten, Jeremy D. Kratz, Madhuchhanda Roy, Stephanie M. McGregor, Jessica D. Lang

**Affiliations:** Center for Precision Medicine, University of Wisconsin-Madison, Madison, WI 53705, USA; Department of Medicine, Division of Hematology, Oncology and Palliative Care, University of Wisconsin-Madison, Madison, WI 53705, USA; University of Wisconsin Carbone Cancer Center, University of Wisconsin-Madison, Madison, WI 53705, USA; William S. Middleton Memorial VA Administration Health System, Madison, WI 53705, USA; Department of Pathology and Laboratory Medicine, University of Wisconsin-Madison, Madison, WI 53705, USA

## Abstract

Patient-derived organoids (PDOs) have transformed translational cancer research, allowing tractable models that better represent clinical features than traditional immortalized cell lines. Here we describe two PDOs with differential responses to carboplatin derived from sequential ascites fluid collections from a patient with high-grade müllerian carcinoma, that could not be further subclassified on the omental biopsy. Uterine origin was clinically excluded by pelvic imaging/CT scan of the uterus and absence of vaginal bleeding. Successful derivation from independent collections enabled comparison of intra-patient heterogeneity across sequential ascites samples and demonstrates that PDO efficiency rate is at least partly patient-specific or tumor-dependent. We performed long-read whole genome sequencing on the two PDOs, OC104 and OC109, to better characterize the structural variant landscape while also obtaining information on single nucleotide variants and DNA methylation. In addition to confirming single nucleotide variants noted in clinical sequencing (TP53, KRAS, SPOP, PPP2R1A, KMT2D), we identified additional variants in TSC2, NCOR2, and CTNNA2 that are predicted to be likely pathogenic. The spectrum of mutations, particularly the coincident KRAS and TP53, highlighted unexpected overlap with ovarian mucinous carcinoma. We also identified larger insertions and deletions that result in non-synonymous variants in MUC5AC, TPRX1, and BMX, as well as four translocation events, including two that could not have been resolved with short-read sequencing. Differentially methylated promoters between the two PDOs include 201 oncogenes and tumor suppressor genes, with HNF1A, MSI2, and SETBP1 having methylation directions consistent with these genes’ roles in platinum response differences observed between the PDOs. Notably, the clonal nature of PDOs produced from two samples taken one week apart is important for the field to appreciate, particularly since they have clonal differences in platinum response. The temporal differences in clonality may indicate a limitation of low volume sampling, however may provide opportunity to longitudinally predict clinical outcomes. We also demonstrate the ability of long-read sequencing to add detail into the genomics and epigenetics of ovarian cancer.

## Introduction

Primary ovarian carcinoma is typically divided into serous, endometrioid, clear cell, and mucinous categories based primarily on morphologic features. While endometrioid and clear cell tumors are both associated with endometriosis and have shared molecular biology, high-grade serous carcinoma is thought to be derived from TP53 mutation-associated transformation of fallopian tube epithelium and the origin of mucinous tumors remains a mystery. Though it lacks an association with endometriosis, serous carcinoma demonstrates discernible phenotypic overlap with endometriosis-associated tumors; mucinous carcinoma is more of an outlier among ovarian cancers with respect to gene expression, though it frequently demonstrates aberrant TP53 and is also associated with hallmarks of low-grade serous tumors, such as KRAS mutation^1–5^. In current practice, the undeniable diagnostic overlap among these entities infrequently affects patient management, because despite such disparate underlying biology, patients with these tumors are generally treated with the same regimen of conventional platinum and taxane-based chemotherapy. Robust models are necessary to explore improved treatment options for these aggressive tumors.

Patient-derived organoids (PDOs) of cancer, which are primary tumor cells typically grown in a dome of extracellular matrix with a specialized cocktail of media rich in supplements and inhibitors of senescence, have been shown to more faithfully recapitulate tumor genomics and therapeutic outcome^6–10^. To date, approximately 178 peer-reviewed studies have been conducted using ovarian cancer PDOs^11^, including five studies with Illumina-based whole genome sequencing on a total of 121 PDOs, predominantly representing high-grade serous ovarian carcinomas^8,9,12–14^. Recently, a mucinous ovarian carcinoma-focused study was posted as a preprint, characterizing 19 long-term PDOs by whole-exome sequencing, RNA-seq, and therapy response for 11 chemotherapy drugs^15^. This study included platinum-based chemotherapy but failed to observe substantial platinum response in PDOs despite an initial patient response. Another report used a library of ovarian cancer organoids of diverse histologic origin, focusing on eleven clear cell carcinoma PDOs^16^. This study performed transcriptome analysis and whole exome sequencing, identifying prominent PIK3CA and ARID1A mutations characteristic of these tumors. They also performed high-throughput drug screens and identified AGR2 involvement in platinum response.

Recent advancements in long-read sequencing (LRS) have made it possible to capture reads exceeding 10 kilobases, greatly enhancing the ability to detect structural rearrangements that are common in ovarian cancer. Oxford Nanopore’s technology has also enabled the capture of DNA methylation from native DNA; that is, without modification typically necessary for bisulfite-conversion-based methods and others^17^. While LRS is more commonly applied in human contexts to RNA to determine splice variants, it has been increasingly used for somatic whole genome sequencing in cancer. Around 70 published studies have investigated long-read sequencing of cancer whole genomes, whether from tumors or cell lines. General themes interrogated in this published body of literature include structural variant detection and genome architecture, extrachromosomal DNA detection, viral integration site identification, DNA methylation, haplotype-specific features, and repetitive elements. Only four studies to date have investigated ovarian cancer whole genomes with the Oxford Nanopore Technologies platform, with three focused exclusively on high-grade serous ovarian carcinomas^18–20^. Sun *et al*. focused on the detection of extrachromosomal DNAs in twelve HGSOC genomes, including focuses on genomic stability and hypomethylation within the ecDNAs^18^. Takamatsu and colleagues sequenced six treatment naive tumors to determine the effect of homologous recombination deficiency on repetitive sequences such as transposable elements, centromeres, and telomeres^19^. Interestingly, they did not identify hypermethylation of BRCA1 or RAD51C in their specimens. Diaz *et al*. interrogated structural variant landscapes of five tumors, primarily focusing on structural rearrangements and association with repetitive sequence^20^. One additional pan-cancer study included nine ovarian tumors, including four clear cell, three high-grade serous, and one each of endometrioid and low-grade serous^21^. Due to the broad nature of this study, ovarian cancer-specific analyses were limited to investigation of homologous recombination deficiency driven by RAD51C allele-specific silencing found in two samples with high homologous recombination deficiency scores. To our knowledge, only three studies have investigated LRS of whole genomes of cancer PDOs, with two studies in esophageal adenocarcinoma focused on extrachromosomal DNAs and de novo assembly of complex rearrangements^22,23^ and an early study that included two breast cancer PDOs that defined structural variants at a high level^24^.

In order to investigate ovarian cancer PDOs using LRS, which is inherently multiomic and better at SV detection, we leveraged two PDOs derived from a single patient. While the original diagnosis could not be specified further than high-grade müllerian carcinoma with p53 abnormality and was therefore regarded as the most common high-grade serous carcinoma, our LRS data demonstrates features overlapping with mucinous ovarian carcinoma, which is unusual as a major diagnostic consideration alongside high-grade serous carcinoma, and of note, more chemoresistant. Interestingly, although the two PDOs are derived from two separate collections only one week apart, they display differential responses to carboplatin. Additionally, some single nucleotide variants, structural variants, and promoter methylation are distinct between the two PDO models, making this patient-matched system quite powerful for understanding potential contributors to treatment resistance in advanced ovarian carcinomas.

## Materials and Methods

### Case information

The patient was identified through routine sample collection. Clinical information was abstracted per manual chart review, including patient age, clinical presentation, pathologic parameters, laboratory results (including Tempus xT sequencing), imaging findings, treatment course, and outcome (UW IRB 2025-0220). Archived diagnostic slides from the initial diagnosis and interval debulking were reviewed and compared by two gynecologic pathologists (SMM, MR).

### Patient-derived organoid development and propagation

Malignant ascites samples were collected through the University of Wisconsin-Madison’s Translational Science Biocore Biobank (UW IRB 2023-1237). The samples used to generate OC104 and OC109 were obtained at the time of aborted interval debulking (after 4 cycles of neoadjuvant chemotherapy) and a subsequent therapeutic paracentesis performed one week after surgery, respectively. Ovarian cancer PDOs were pelleted, Ficoll-separated (Cytiva, Uppsala, Sweden), and resuspended in base media prior to embedding at a 1:1 ratio in Cultrex Growth Factor Reduced Basement Membrane Extract (R&D Systems, Minneapolis, MN, USA). For at least two passages, organoid development media (**Supplemental Table 1**) was used, and for subsequent long-term culture and experiments, long-term propagation and experiment media (**Supplemental Table 1**) was used.

### Histochemical staining of patient-derived organoids

Cell pellets were processed into formalin-fixed, paraffin embedded blocks and was performed per routine methods by the UW-Madison Translational Research Initiatives in Pathology (TRIP), which also performed hematoxylin and eosin (H&E) staining. H&E slides of PDOs were reviewed by two gynecologic pathologists involved in this study (MR & SMM).

### Growth measurements of patient-derived organoids

PDOs were plated in 12-well plates as 100 µL hanging-drop domes and imaged at 0 h and 96 h using the 4x objective of the Cytation5 (Agilent, Inc.) with a 5-field Z-stack (±2 fields of view x250 µm from the autofocused plane) and 5×5 field montage to create a full field of view.

Subsequent images were analyzed in Gen5 3.17 and underwent image preprocessing, stitching, and z-projection to control for background noise, produce a larger field of view, and render a z-stack, respectively. Individual objects were defined using an automated contrast threshold and included only when object diameter fell between 50 µm and 750 µm. Each object was assessed for diameter, area, x-coordinate, y-coordinate, and circularity. Object metrics were imported into MATLAB version: 23.2.0 (R2023b) to pair corresponding datapoints across the time course, using circularity >0.4 and Euclidean distance <75 µm as mapping criteria using the Statistics and Machine Learning Toolbox. The normalized change in area for each object was calculated between 0 h and 96 h.

### Flow cytometry DNA content assay

PDOs were harvested and broken up into a single cell suspension using TrypLE (Gibco).

CAL51 cells were harvested with 0.05% Trypsin. After washing with PBS, 1 million cells were aliquoted into a separate tube and two drops of Hoechst 33342 Ready Flow™ Reagent (Invitrogen) were added to the cells. Cells were incubated at 37°C for 1 hour, with gentle vortexing after 30 minutes to ensure even staining. After staining, cells were spun down and resuspended in 500μL PBS and analyzed on a Cytoflex instrument. Data were analyzed using FlowJo.

### Drug dose response assay

Drug dose response assays for carboplatin were performed on PDOs OC104 and OC109. PDOs were plated in 10μL of 1:1 mixture of base medium (Advanced DMEM/F12 medium + 1% penicillin/streptomycin + 1% GlutaMAX) and Cultrex in a μ-Plate 96 well 3D plate (Ibidi, Fitchburg, WI) at about 4,000 small organoids/well. Fifty μL of long-term propagation and experiment media was added to the well after polymerization of Cultrex, and PDOs were further incubated for 24 hours. Culture medium was replaced with 60 μL carboplatin-containing complete organoid medium (carboplatin from Cayman Chemical Company, cat.13112). Each PDO was tested at 12-doses that ranged between 1mM to 500nM with serial 1:2 dilutions.

Carboplatin treatment continued for 72 hours, followed by replacement with fresh complete organoid medium for an additional 72 hours. Media was removed and 20 μL of CellTiter-Glo 3D (Promega) was added to each well and incubated according to the manufacturer’s instructions. Luminescence was read using a BioTek Cytation 5 Cell Imaging Multi-mode Reader (Agilent). Background-subtracted luminescence readings were analyzed in GraphPad Prism 9 with non-linear regression fit to calculate an IC_50_ value. The mean of seven technical replicates was calculated for each of two independent experiments, and the means of each independent experiment were used to calculate the IC_50_ value. Drug dose response assays for each PDO were performed at the same time since patient sample acquisition.

### Long-read sequencing preparation

Genomic DNA was isolated from cells using a Quick-DNA Miniprep Plus Kit (Zymo Research). Sequencing libraries were prepared according to Oxford Nanopore’s manufacturer’s protocol for the ligation sequencing kit V14. Total run time for the sequencing was set to 100hrs. Flow cells were washed and re-loaded after 48hrs. The target yield for each sample was 120GB.

### Long-read sequencing analysis

Base calling was performed on the sequencer using Dorado (MinKNOW v23.07.12; Oxford Nanopore Technologies) using the High-accuracy model, 400 bps model (dna_r10.4.1_e8.2_400bps_sup@v4.2.0). Reads were aligned to a version of the hg38 reference genome containing only chromosomes 1-22 and X. The EPI2ME wf-human-variation pipeline (v2.6.0; github.com/epi2me-labs/wf-human-variation) was used to identify structural variants in each of the two PDOs.

Single nucleotide polymorphisms (SNPs) were called within the EPI2ME wf-human-variation pipeline. VCF files were parsed to filter for MANE Select protein-coding variants with a Consequence of stop_gained, frameshift_variant, splice_acceptor_variant, splice_donor_variant, or missense_variant. Missense variants were required to be called as damaging/pathogenic by at least two of 3 of SIFT, PolyPhen, or AlphaMissense variant effect predictors, as annotated with VEP. Variants with allele fractions (AF) in ExAC, the National Heart Lung and Blood Institute GO Exome Sequencing Project, or the Thousand Genomes Project cohorts greater than 0.01 were excluded, and only COSMIC-annotated oncogenes and tumor-suppressor genes were prioritized. Variants were manually inspected in Integrative Genomics Viewer, for which we also imposed a requirement of a QUAL score of >20, which removed all poorly supported variant calls, as well as independent confirmation of AF in GnomAD less than 0.01.

Copy number analysis was performed using CNVkit (v0.9.13) to call bin- and segment-level copy number ratios from bam files. CNVkit was also used to plot copy number figures.

Structural variants were then called using Sniffles2 (v2.0.7-epi2me)^25^ and CuteSV (v2.1.1)^26^. MAVIS (v3.1.2)^27^ was used to merge and annotate SVs. Because we did not have germline samples for the cell lines, and because many of the filtered SV sites had low coverage in the gnomAD (v3)^28^, we further annotated mean proportion of individuals with coverage greater than 20x (referred to as >20x) and greater than 50x (referred to as >50x) across each SV region, or at both breakpoint sites of translocations. SVs with population fraction with coverage >20x of more than 0.9 and population fraction with coverage >50x of less than 0.05 were considered to have a low likelihood of being normal population variants; these were considered for further analysis, with the exception of translocations, where the coverage at the two break points was often discrepant and infrequently associated with other common features of common genome structural variation (*i.e.*, repetitive sequences). Translocations were visually inspected in Integrative Genomics Viewer (v2.18.1). Translocations were excluded from further analysis if they were found in our high-grade serous ovarian carcinoma LRS datasets (highly indicative of poor mapping or common variants; data forthcoming in another publication), if other reads at the site of the translocation supported an insertion with read-through on the right and left ends of the region mapping to the alternate chromosome, or if reads poorly supported the translocation call on visual inspection. Structural rearrangements were plotted as ideograms using CyDAS.

Methylation was from long-read sequencing data. Haplotype-phased bedMethyl outputs from the EPI2ME human-variation pipeline were merged using modkit (v0.6.1; ONT) to generate sample-level bedMethyl files for 5-methyl-cytosine. Differentially methylated regions (DMRs) were called using Dispersion Shrinkage for Sequencing data (DSS; v3.2)^29^. Only methylation sites with coverage ≥ 5 were used for analysis. Differentially methylated loci (DMLs) were called using DMLtest with the following options: equal.disp = FALSE, smoothing = TRUE, smoothing.span = 500. DMLs were used to call DMRs using callDMR with the following options: p.threshold = 1e-4, minlen = 100, minCG = 15, dis.merge = 100, and delta = 0. DMRs were annotated to promoters using annotations from the TxDb.Hsapiens.UCSC.hg38.knownGene package in R. Oncogenes and tumor suppressor genes were annotated from the Catalogue of Somatic Mutations in Cancer (COSMIC) reference dataset downloaded 12/02/2022 with Tier 1 evidence. Manhattan plots were plotted using plotgardener.

### Variant analysis of AACR Project GENIE

Project GENIE data (v19.0-public) from the American Association for Cancer Research (AACR) was accessed through the cBioPortal interface. “Ovarian Cancer” was selected in the Cancer Type field. Only subtypes currently accepted by the WHO Classification of Tumours “Female Genital Tumours” 5th edition were included in the analysis. Each gene was queried for each histotype separately with the option “show only profiled > in all queried genes” option.

Specific mutations were searched in the “Mutations” tab, and the number of patients with the mutation was divided by the total number of patients profiled to yield the specific mutation’s frequency.

### Quantitative, Real-Time PCR

Total RNA was isolated from organoid samples (OC104 and OC109) with Zymo Quick-DNA/RNA MiniPrep Plus Kit (Cat#7003, Zymo Research Corporation, Irvine, California, USA). Complementary DNA (cDNA) was generated from purified RNA using Bio-Rad iScript cDNA Synthesis Kit (Cat# 1708891, Bio-Rad, Hercules, California, USA) using 700ng of RNA. The 20 μL reaction volume was diluted with 80 μL nuclease-free water prior to qRT-PCR. Gene expression was assessed using quantitative reverse transcription polymerase chain reaction (qRT-PCR). Primer sets were designed for the genes HNF1A, MSI2, MDM4, SETBP1, BRCA2 and the housekeeping gene RPLP0. Primer sequences are listed below. Oligonucleotide primers were synthesized by IDT (Integrated DNA Technologies, Coralville, Iowa, USA). qRT-PCR reactions were performed using Bio-Rad iQ SYBR Green Supermix (Cat# 1708880, Bio-Rad, Hercules, California, USA) in a total reaction volume of 10 μL containing 5 μL iQ SYBR Green Supermix, 500 nM forward primer, 500 nM reverse primer, and 2 μL cDNA template. A no-template control (NTC) was included in each assay to monitor for contamination and nonspecific amplification. All reactions were performed in triplicates. Amplification was performed on the QuantStudio 6 Pro using default cycling conditions with melt curve. Relative gene expression levels were normalized to RPLP0 and calculated using the comparative Ct (2^−ΔΔCt) method.

Primers: HNF1A Forward: 5’-CTCTCCCCCAGTAAGGTCCA-3’; HNF1A Reverse: 5’-TGAGACCAGCTTGGCTTCTG-3’; MSI2 Forward: 5’-GCACAGAGGGTTTGGCTTTG-3’; MSI2 Reverse: 5’-TGAACGCGTCCATGGTGTAA-3’; MDM4 Forward: 5’-AGGCCCTAGGATCTGTGACT-3’; MDM4 Reverse: 5’-CCAGGAGAGATCCTGCAAGC-3’; SETBP1 Forward: 5’-AGACCACAAAGCGGGCTAAG-3’; SETBP1 Reverse: 5’-CAAGGTCACTGGCTGTGAGG-3’; BRCA2 Forward: 5’-TGCACCTCTGGAGCGGA-3’; BRCA2 Reverse: 5’-AGGTTCAGAATTATAGGGTGGAGC-3’; RPLP0 Forward: 5’-GACAATGGCAGCATCTACAAC-3’; RPLP0 Reverse: 5’-GCAGACAGACACTGGCAAC-3’.

## Results

### Summary of case

The 68-year-old patient with no known relevant medical history or smoking history presented at the ER with a one-month history of bloating, abdominal distension, decreased appetite, fatigue, and constipation. CA125 was elevated at 255 U/mL and abdominopelvic CT demonstrated a 7.7 x 6.8 x 5.4 cm left pelvic mass, with ascites and peritoneal implants. The patient underwent a core needle biopsy of the omentum. The diagnosis was high-grade müllerian carcinoma, supported by marked nuclear atypia and immunophenotypic findings including PAX-8 positivity and a null pattern for p53 (**Figure 1A**). Additional findings included negativity for WT1 and estrogen receptor alpha. More specific classification was not possible, but the findings were compatible with either high-grade serous carcinoma of tubo-ovarian origin or uterine serous carcinoma, with the former being the most likely scenario based on disease spread and incidence in routine clinical practice. Clinically, there was no history of vaginal bleeding, and the uterus was visualized by CT with no indication of abnormality, supporting ovarian origin, for a working diagnosis of at least stage IIIC tubo-ovarian high-grade serous carcinoma. Accordingly, the patient underwent 4 cycles of neoadjuvant chemotherapy with carboplatin/paclitaxel/bevacizumab; by imaging, there was reduction in size of the ovarian mass but increased volume of ascites and possible increase in peritoneal carcinomatosis. Interval debulking was attempted, but the surgery was aborted due to extensive intraabdominal disease involving peritoneal surfaces, bowel, and lesser omentum/porta hepatis. Resection of disease would have required extensive peritoneal resection, extensive bowel resection (possibly total colectomy with permanent ostomy), and radical upper abdominal surgery given lesser sac and porta hepatis disease. With likely inability to achieve optimal resection status even with these radical procedures, a decision was made to abort the procedure in favor of additional chemotherapy. Post-treatment figures demonstrating some response shown in **Figures 1B-C**. Ascites fluid was collected during the aborted surgery, which was used for the development of OC104. Limited molecular characterization was performed to identify possible treatment options, revealing KRAS G13D, SPOP E50K, PPP2R1A S256F, TP53 E294*, and KMT2D Q3507*. The tumor was microsatellite stable and negative for homologous recombination deficiency, with TMB of 4.2 m/MB. In addition to the clinical sequencing variants, FOLR1 was negative by immunohistochemistry. Due to persistent high-volume ascites, weekly paracentesis was performed; OC109 was developed from a repeat sample obtained from the first post-surgical therapeutic paracentesis performed at an interval of one week. After one additional cycle of carboplatin/paclitaxel/bevacizumab with progressive disease by imaging, the drug regimen was transitioned to liposomal doxorubicin and bevacizumab. The patient developed complications and ultimately died of disease just over four months after attempted surgery.

**Figure 1.**
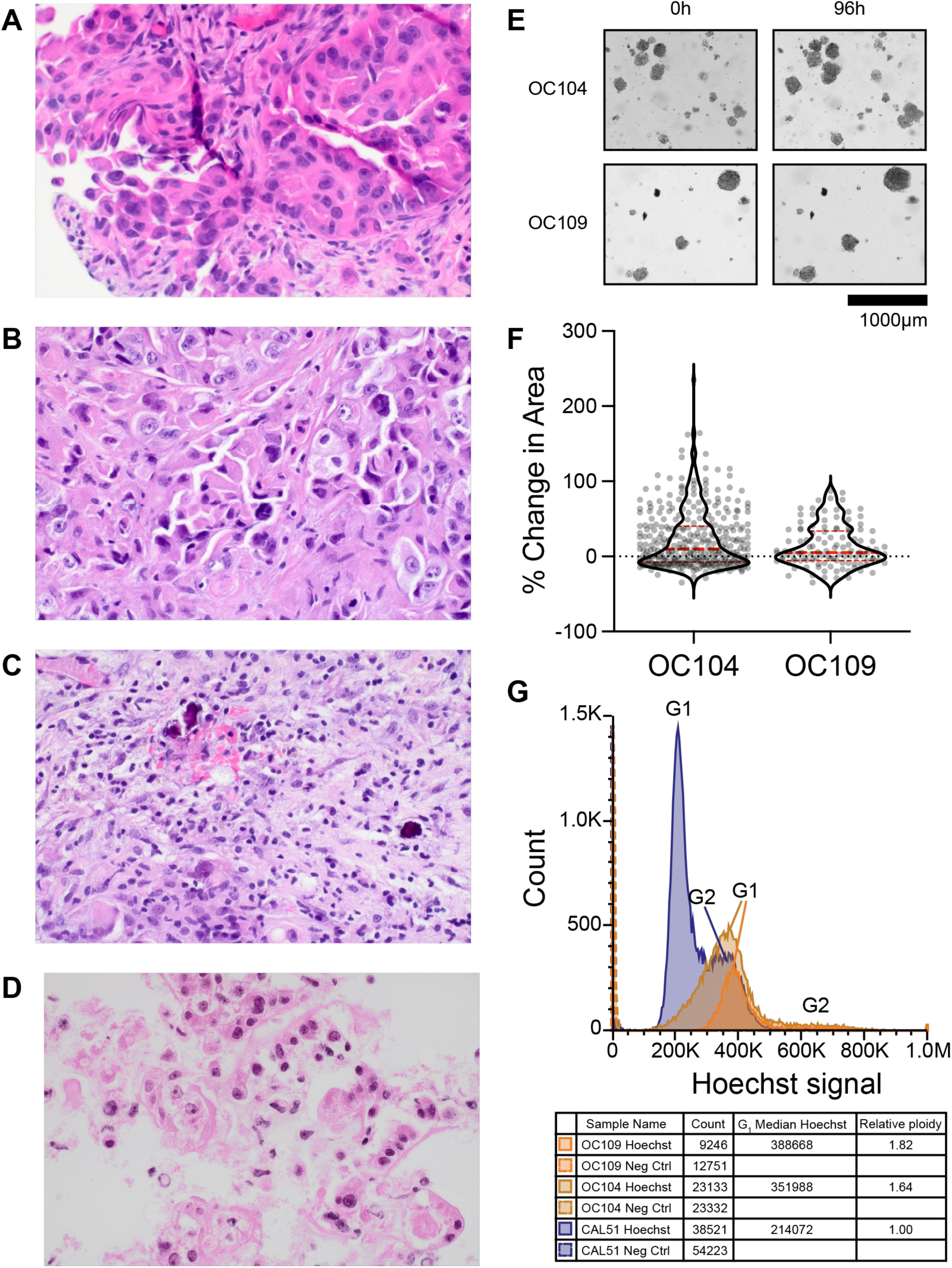
Tumor and patient-derived organoid OC104 and OC109 characteristics. (A) The omental biopsy demonstrates high grade morphology including large and markedly atypical nuclei and mitosis indicated by red circle (hematoxylin-eosin, original magnifications x 400). (B) Post-treatment peritoneal biopsy demonstrating residual tumor with high grade morphology including large and markedly atypical nuclei and mitosis (hematoxylin-eosin, original magnifications x 400). (C) Partial tumor response is evident by fibrosis associated with macrophages, mixed inflammatory cells, and psammoma bodies (hematoxylin-eosin, original magnifications x 400). (D) PDO morphology similar to the patient’s tumor. (E) Representative microscopy images of each PDO model over 96 hours of culture. (F) Changes in individual organoid areas change at 96h relative to 0h. Red indicates mean change in organoid area. OC104 showed a mean increase of 22.41% (± 39.84 SD) and OC109 showed 14.20% (± 27.54 SD). Number of individual measured organoids per PDO: OC104 N = 334, OC109 N = 116. (G) DNA content analysis of PDOs vs. diploid cell line, CAL51. Cells were gated for single cells based on forward/side scatter (data not shown), then plotted on linear scale as a histogram. Count indicates how many single cells were captured. G1 median Hoechst indicates the median value of Hoechst signal for the G1 peak for each experimental value. Relative ploidy is the fold difference in value of PDOs compared to the diploid cell line control. DNA content analysis was performed approximately 9 months after ascites collection.

### Development of patient-derived organoids

Ascites collections were obtained at two times, one week apart from each other, to generate PDOs OC104 and OC109 (**Figure 1D**). PDO growth was assessed within 43-50 days of sample acquisition for a period of 96 hours by measuring changes in organoid area. Both lines exhibited a net positive mean change in area, with OC104 showing a mean increase of 22.41% (± 39.84 SD) and OC109 showing 14.20% (± 27.54 SD) (**Figure 1E-F**). The greater standard deviation in OC104 reflects higher inter-organoid variability compared to OC109. A subset of organoids in both lines exhibited negative area changes, consistent with the biological heterogeneity characteristic of patient-derived cultures.

To support the malignant origin of the organoid cultures, we assessed DNA content and observed aneuploidy consistent with genome doubling using a DNA content assay, representing a 1.64- and 1.82-fold increase in total DNA content relative to diploid controls for OC104 and OC109, respectively (**Figure 1G**).

### Confirmation of clinical sequencing variants in patient-derived organoids with long-read sequencing and novel potentially pathogenic variants

As mentioned above, targeted clinical sequencing using Tempus xT comprehensive genomic profiling revealed that the tumor bears pathogenic mutations to KRAS (G13D), SPOP (E50K), PPP2R1A (S256F), TP53 (E294*), and KMT2D (Q3507*) (**Table 1**). The tumor was microsatellite stable, negative for homologous recombination deficiency, and had a tumor mutation burden of 4.2 m/MB. LRS was performed on OC104 and OC109, which confirmed all variants identified in clinical genomics sequencing (**Table 1**). While allele fractions were overall higher in OC104 and OC109 in comparison to the clinical genomics report, this is likely due to a higher tumor purity in PDOs. The highest AFs were 0.8966 and 0.9756 for SPOP in OC104 and OC109, respectively, indicating >90% pure cancer organoids. In addition to the five variants identified from the clinical report, we also prioritized missense variants in NCOR2, TSC2, and CTNNA2 for further investigation based on our prioritization strategy (**Table 1** and **Supplemental Table 2**). Most of these mutations were found at allele fractions less than 0.5, suggesting subclonal variants or copy number gains in these genomic regions. NCOR2 is only found in OC104, indicating a clonal variant. IGV validation snapshots for the non-clinical variants are shown in **Supplemental Figure 1**.

**Table 1.** Single nucleotide variants reported in a clinical sequencing report from tumor sequencing at Tempus. Allele Fraction (AF) is reported from the clinical sequencing report (Clinical Genomics AF), as well as from long-read sequencing of OC104 and OC109 PDOs. “na” indicates that the variant was not detected/reported.

NCOR2, Nuclear receptor corepressor 2, had a p.Arg1625Cys characterized by PolyPhen to be “probably damaging” and by SIFT to “deleterious (low confidence)” (**Supplemental Table 2**). Arginine 1625 is not in an annotated protein domain of NCOR2, but the AlphaFold model places this residue between the two DNA binding domains and the SMC N-terminal domain (**Figure 2**). This variant has not been observed in ovarian cancers in the GENIE database (**Supplemental Table 2**).

**Figure 2.**
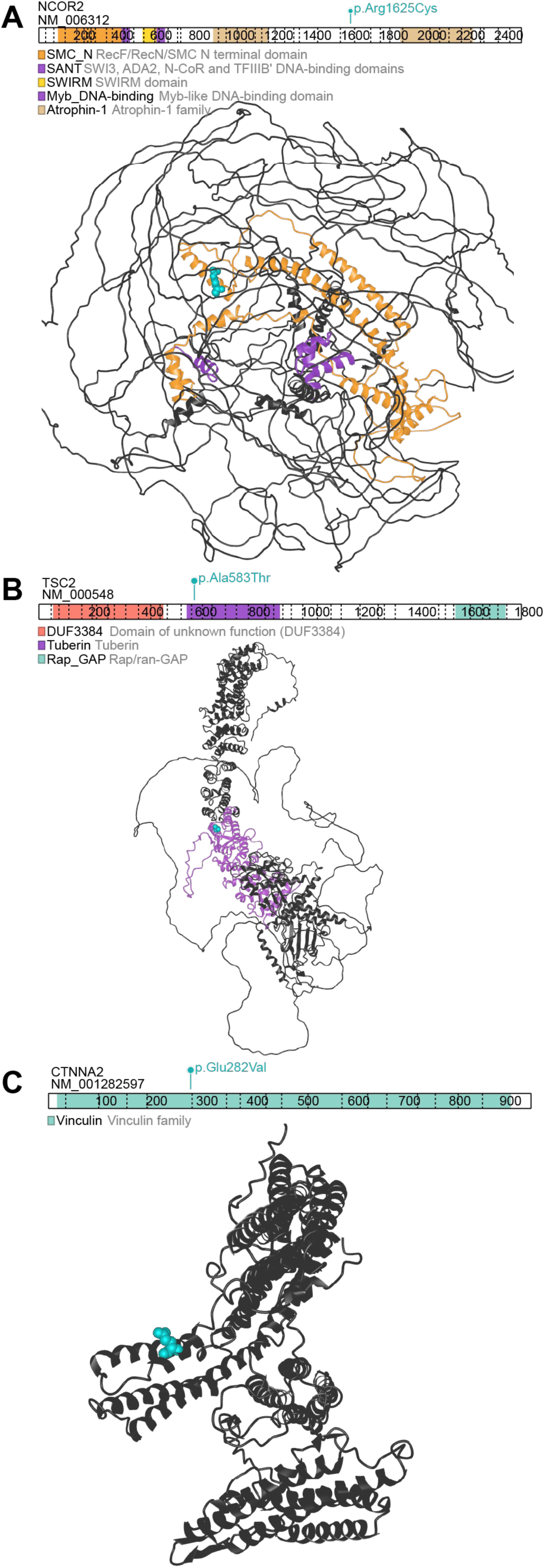
Novel potentially pathogenic missense mutations identified by long-read sequencing. (A) NCOR2 domain structure highlighting the p.Arg1625Cys missense variant and AlphaFold predicted structure (Q9Y618-4). The p.Arg1625 position is shown in cyan. SMC domain is shown in orange, and DNA binding domains are shown in purple. (B) TSC2 domain structure highlighting the p.Ala583Thr missense variant and AlphaFold predicted structure (P49815-5). The p.Ala583 position is shown in cyan and the Tuberin domain is shown in purple. (C) CTNNA2 domain structure highlighting the p.Glu282Val missense variant and AlphaFold predicted structure (P26232-2). The p.Glu282 position is shown in cyan.

TSC2, which encodes tuberin, had a p.Ala583Thr missense variant characterized by PolyPhen to be “probably damaging” and by SIFT to “deleterious (low confidence)” (**Supplemental Table 2**). Alanine 583 is located within the tuberin domain of the protein, specifically near the end of an alpha helix towards the center of the protein (**Figure 2B**). This missense mutation has been observed in the GENIE cohort of ovarian cancers, specifically in high-grade serous ovarian carcinomas and endometrioid ovarian cancers, as well as unspecified ovarian epithelial cancers (**Supplemental Table 2**).

CTNNA2, which encodes alpha-catenin, had a p.Glu282Val variant characterized by AlphaMissense to be “likely pathogenic” and by SIFT to be “deleterious (low confidence)” (**Supplemental Table 2**). Glutamic acid 282 is within the vinculin domain of the highly structured alpha-catenin protein structure, and sits at the MI four-helix bundle that binds to vinculin and mediates cell-cell adhesion^30^ (**Figure 2C**). CTNNA2 is not included in the gene panels used in GENIE.

Given that the concurrent presence of high allele fraction KRAS and TP53 mutations is rare for high-grade serous ovarian carcinoma, we interrogated a large somatic sequencing dataset, AACR’s GENIE Project, to determine which ovarian cancer subtypes best match the molecular profile of our PDOs. Both those SNPs identified in clinical sequencing and from our LRS were interrogated, using both exact mutation matches (**Figure 3A**) and any mutation to the gene (**Figure 3B**). CTNNA2 was not sequenced as part of this effort and was excluded from analysis. Exact mutation matches were overall very low, representing fewer than 4% of cases for any subtype/gene (**Figure 3A**). While TP53 mutations were the strongest matches, it was nonspecific for determining subtype. The sum of altered cases across genes was greatest for mucinous ovarian cancer and endometrioid ovarian cancers. High-grade serous ovarian carcinoma similarity is largely driven by TP53 presence only.

**Figure 3.**
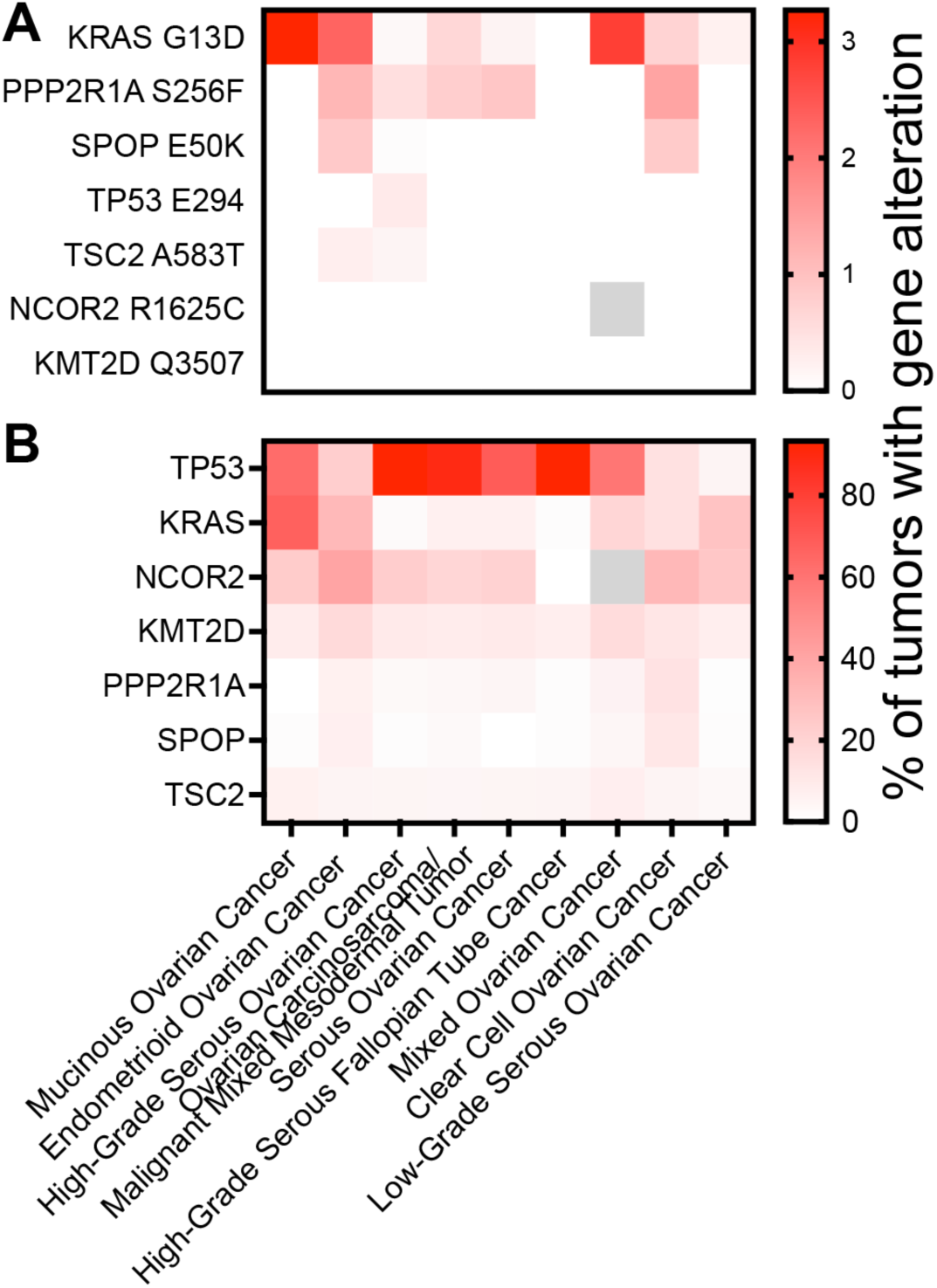
Frequency of genetic alterations to the genes altered in OC104/OC109 in ovarian cancers from the AACR Project GENIE database by molecular subtype. (A) Frequency of exact mutations found in OC104/OC109 by subtype. (B) Frequency of any mutation in the genes altered in OC104/OC109 by subtype.

### Structural variant findings in long-read sequencing

Long-read sequencing was performed on the two PDOs to interrogate the structural variants, since the original diagnosis as high-grade müllerian carcinoma might be suggestive of high-grade serous ovarian carcinoma. High-grade serous ovarian carcinomas bear a high copy number burden and many structural rearrangements. Unfortunately, germline (blood) samples were not acquired from the biobank for this patient. Sequencing was performed on an Oxford Nanopore Technologies PromethION24, and EPI2ME’s wf-human-variation pipeline was used for germline analysis.

PDO copy number analysis revealed that overall copy number profiles were similar between OC104 and OC109, though some heterogeneity exists between them (**Supplemental Figure 3A and B**). Largest copy number gains were at 8q and 12p, while copy number loss was observed at 9p. 9p contains the CDKN2A locus, which shows copy number loss in both OC104 and OC109 (**Supplemental Figure 3C**).

Structural variants (SVs) were called using sniffles2 and cuteSV, then merged using MAVIS, which outputs a merged set of SVs, a filtered set of SVs, and a summary of nonsynonymous SVs (**Figure 4A**). After visual inspection of filtered SVs from MAVIS, we identified that the vast majority of SVs were common in high-grade serous ovarian carcinoma cell lines (subject of separate manuscript), likely highlighting population germline variants that still passed GnomAD filters. Additionally, many of the regions containing SVs were outliers in their population-based coverage, based on the public germline sequencing database, gnomAD. To address this, we added a filter focusing on SVs from regions in which a high fraction of the population had coverage near the genome-wide mean coverage (**Figure 4B** and **Supplemental Figure 4**). The rationale is that regions in which a large fraction of the population deviates substantially from the genome-wide mean coverage are likely to have common copy number or structural variation. This step mainly reduced the number of insertions and deletions.

**Figure 4.**
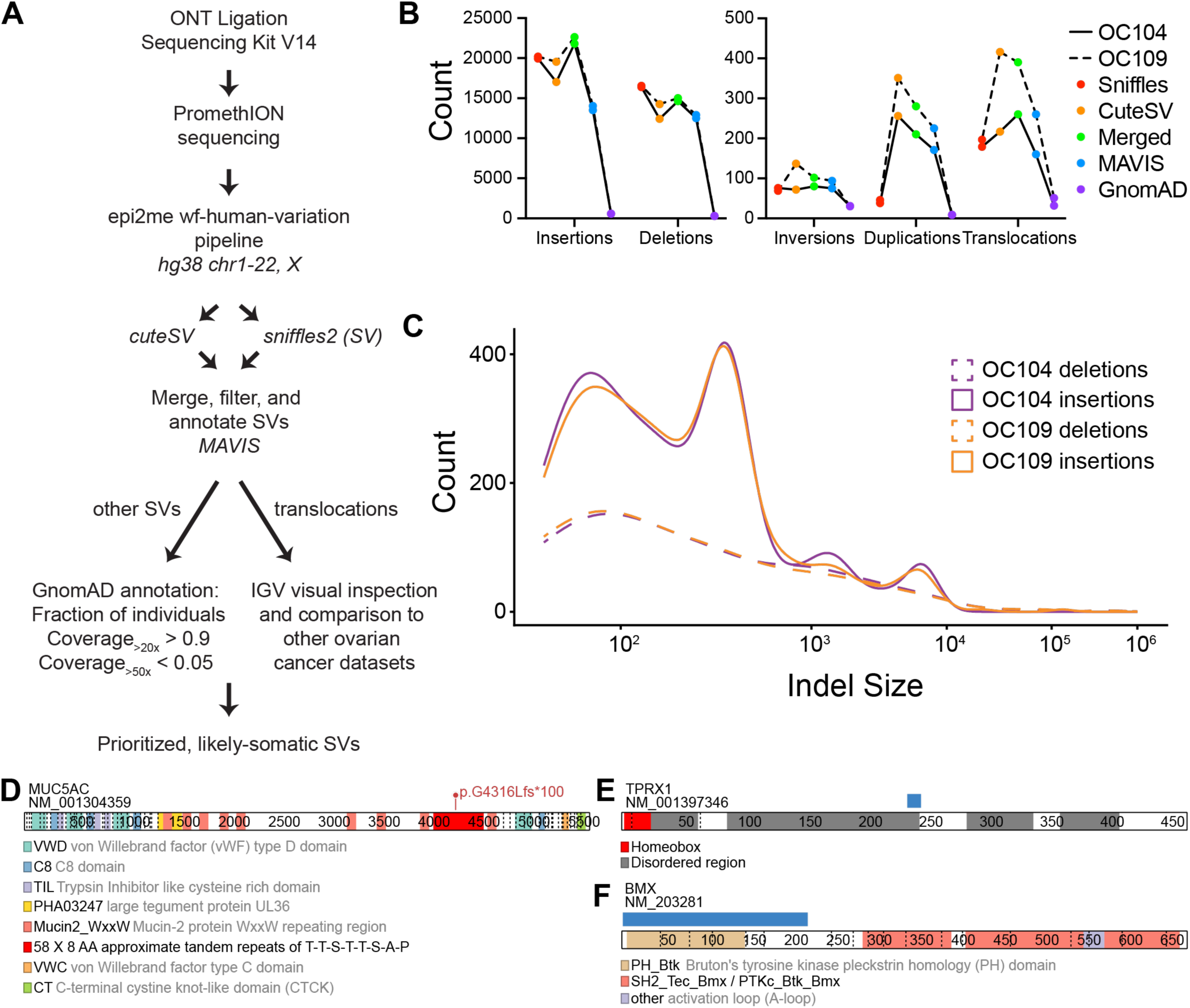
Long-read sequencing structural variant summary and nonsynonymous structural variants. (A) Workflow of ONT LRS pipeline and analysis steps. (B) Number of each SV event type by stage in the filtering pipeline. (C) Distribution of lengths of insertions and deletions (Indels) in base pairs. (D) MUC5AC protein domain map with predicted protein consequence (p.G4316fs*100) resulting from a deletion. Annotated domains are highlighted and described in the figure key. Numbers indicate amino acid position. (E) TPRX1 protein domain map with predicted protein consequence (p.G237_P248delGPIPGPISGPNPinsA) resulting from a deletion. Deletion is shown in the blue rectangle. Annotated domains are highlighted and described in the figure key. Numbers indicate amino acid position. (F) BMX protein domain map with predicted protein consequence (p.D2_I223delDTKSILEELLLKRSQQKKKMSPNNYKERLFVLTKTNLSYYEYDKMKRGSRKGSIE IKKIRCVEKVNLEEQTPVERQYPFQIVYKDGLLYVYASNEESRSQWLKALQKEIRGNPHLLVKYHSGFFVDGKFLCCQQSCKAAPGCTLWEAYANLHTAVNEEKHRVPTFPDRVLKIPRAVPVLKMD APSSSTTLAQYDNESKKNYGSQPPSSSTSLAQYDSNSKKIinsTTNQRKT) resulting from a deletion. Deletion is shown in the blue rectangle. Annotated domains are highlighted and described in the figure key. Numbers indicate amino acid position.

Independent analysis demonstrated that the peak in insertions and deletions found at 300bp is predominantly driven by repetitive elements such as Alu repeats (**Supplemental Figure 4** and supporting data in forthcoming manuscript), which provides further support for the use of the gnomAD filter to remove common population SVs. In total, the mean numbers of SVs called using this workflow were 583 insertions, 305 deletions, 31 inversions, 9 duplications, and 42 translocations. The identified SVs were then individually inspected in IGV. The size of indels was biased towards smaller indels, with the exception of some predictable, likely repetitive-element-derived insertion peaks (**Figure 4C**).

No SV that passed our filtering and inspection criteria altered protein coding sequence. If the GnomAD coverage filter is loosened, 3 nonsynonymous deletions are observed in OC104 and OC109 that are not also observed in the vast majority of LRS datasets during IGV validation (high-grade serous ovarian carcinoma cell lines; subject of parallel manuscript). The MUC5AC deletion results in a frameshift alteration resulting from a 169 bp deletion on one allele (**Figure 4D**). It is present in both OC104 and OC109. This deletion is contained in a region with over 50 repeats of the 8-amino-acid repeat T-T-S-T-T-S-A-P. Deletions encompassing this region are reported in the GnomAD database, with the most common overlapping variant at an AF of 0.00285 (11:1190869-1191178). MUC5AC has been implicated as a biomarker in mucinous ovarian carcinoma^31–34^ and is predicted to be a target of nonsense-mediated decay based on its position within the transcript. However, we were unable to design a targeted assay to assess allele-specific expression due to the lack of SNPs within exons in the MUC5AC transcript, and we were unable to design specific primers spanning the deletion site.

A homozygous 36 bp deletion within TPRX1 leads to an in-frame replacement of 12 amino acids from G237-P248 with an alanine and was identified in both PDOs (**Figure 4E**). This variant was biallelic in both PDOs. TPRX1 encodes a transcription factor important in the maternal to zygotic transition in early development. The DNA binding domain is the only well characterized domain, and lies far upstream of the deletion. This exact deletion is not reported in GnomAD, and the combined sum of overlapping deletions have an allele fraction of 0.000525 with no homozygotes, indicating that it is quite infrequent.

Deletion of 66 bp in the BMX gene leads to replacement of amino acids 2-223 with T-T-N-Q-R-K-T on one allele in OC109 only (**Figure 4F**). The only deletion in this region in GnomAD is a 28.9 Mb deletion reported from one individual. The deleted region, which is contained within exon 7 and encodes the Bruton’s tyrosine kinase (BTK) pleckstrin homology (PH) domain, is not found in crystal structures of BMX, but is predicted with high-confidence by AlphaFold to have some beta sheet structure. Comparatively, BMX has reported alternative splice variants leading to deletion of exons 1 through 8, which has been implicated in lung cancer^35^.

Merged translocation calls from two LRS SV callers were visually inspected in IGV. From 89 translocations in each of OC104 and OC109 samples, four translocation events appeared to be reasonable translocation calls that are not identified broadly in other LRS samples (high-grade serous ovarian carcinoma cell lines; subject of parallel manuscript). The first results in a dicentric chromosome from a translocation between chromosomes 19p and Xp (**Figure 5A and Supplemental Figure 5A**). The X chromosome was also involved with two other translocation calls: between Xp and 17p near the centromeres (**Figure 5B and Supplemental Figure 5B**), and a complex rearrangement of two sites on X with 6q and 21q that only occurs in OC109 (**Figure 5C and Supplemental Figure 5C**). An inverted translocation also leads to replacement of part of the 2p arm with 21q (**Figure 5D and Supplemental Figure 5D**).

**Figure 5.**
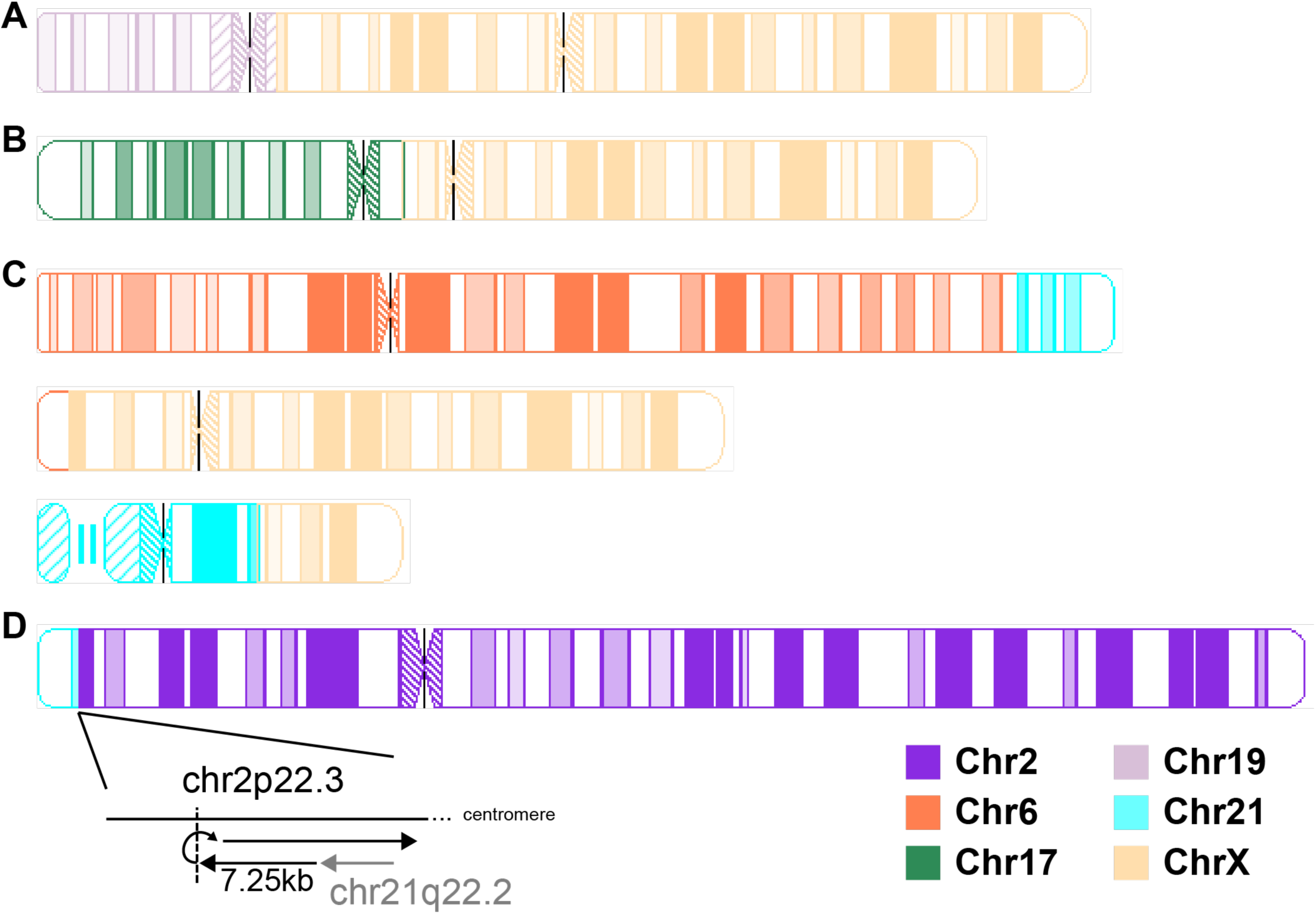
Ideograms of LRS translocation calls from PDOs. (A) BND1, found in OC104 and OC109. dic(X;19)(p22.32;p12). (B) BND2, found in OC104 and OC109. dic(17;X)(p11.2;p11.23). (C) BND3, found only in OC109. der(6)t(6;21)(q27;q21.3), der(X)t(6;X)(q27;p21.1), and der(21)t(21;X)(q21.3;q26.1). (D) BND4, found in OC104 and OC109. der(2)t(2;21)(p22.3;q22.2). Black lines and constrictions indicate centromeres.

### Differentially methylated regions between PDOs

DNA methylation data is also discerned using ONT LRS based on differential voltage of modified vs. unmodified cytosine nucleotides. We specifically examined differentially methylated regions (DMR) between OC104 and OC109 by focusing on 5-methyl-cytosine modifications. A total of 111,345 DMRs were identified between the two PDOs; 105,603 with more methylation in OC104 and 5,742 with more methylation in OC109 (**Figure 6A**). After annotation to genes, 17,904 DMRs overlapped promoters; 15,656 with more methylation in OC104 and 2,248 with more methylation in OC109 (**Figure 6B**). DMR promoters were associated with 201 known oncogenes and tumor suppressor genes (TSGs). We further filtered DMRs by considering a region to be differentially methylated if the absolute difference in methylation proportion between the two samples was ≥L0.4. Forty-nine oncogenes and TSGs that passed this filter are shown in **Figure 6C**.

**Figure 6.**
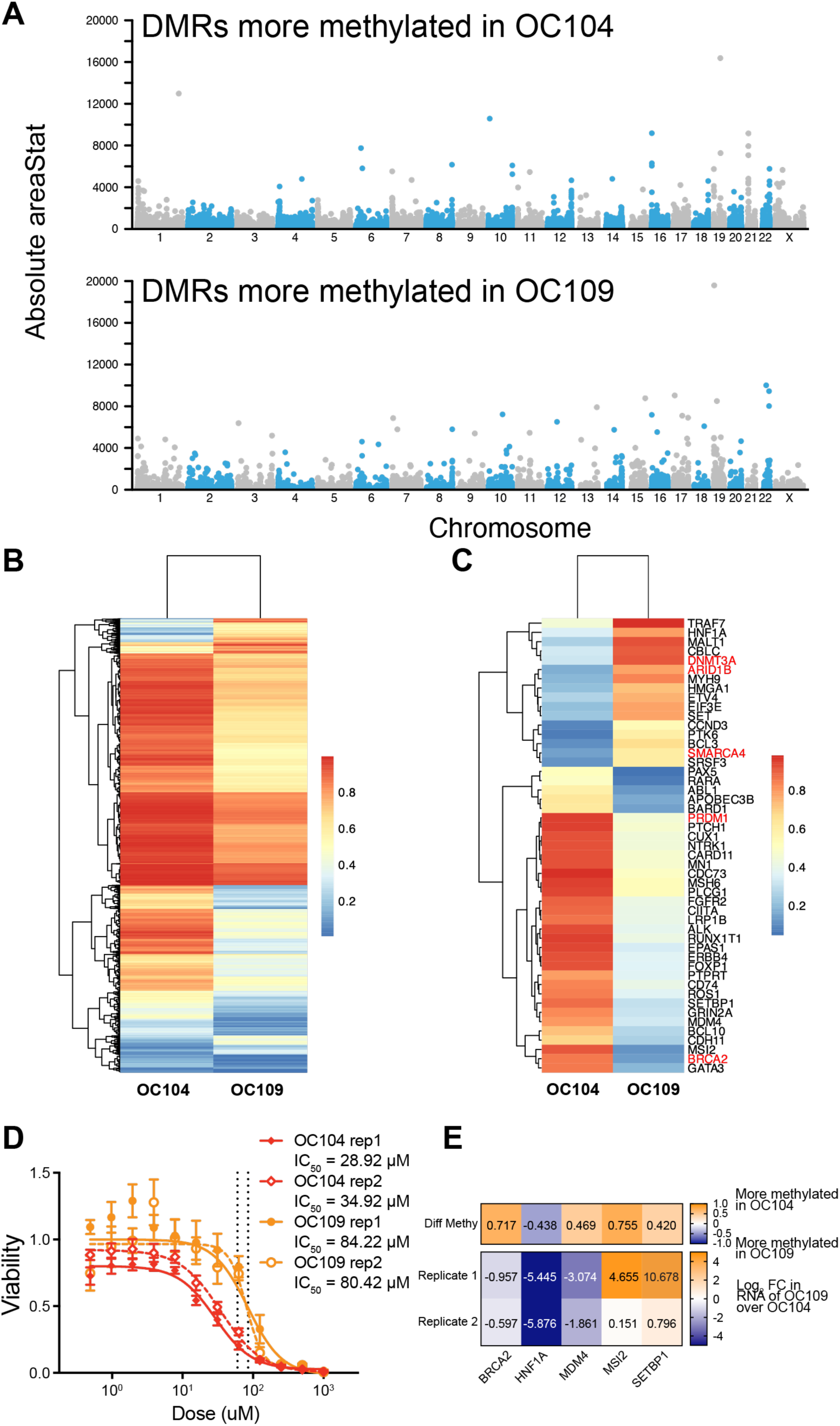
Differential methylation and carboplatin response in PDOs. (A) Manhattan plot of all DMRs identified between OC104 and OC109. Those with positive diff.Methy values indicated higher methylation in OC104 and are shown in the top panel, and those with negative diff.Methy values indicate more methylation in OC109 and are shown in the bottom panel. AreaStat is a measure combining the magnitude of difference in methylation between PDOs and the size of the DMR. (B) Heatmap of all promoter-annotated DMRs in OC104 and OC109. A value of 1 indicates full methylation of DMR, and a value of 0 indicates a fully unmethylated DMR. (C) Heatmap presented as in panel (B), but only showing oncogenes and tumor suppressor genes. Gene names highlighted in red are those later prioritized for RNA expression analysis in panel E. (D) Carboplatin drug dose response assay to determine IC50. Two independent experiments were performed (rep1 and rep2), each with 7 technical replicates per experiment. The maximum clinically achievable dose (Cmax) range of carboplatin is shown by the dotted vertical lines. Points indicate the mean of technical replicates with standard error. (E) qRT-PCR results on RNA extracted from OC104 and OC109 in two replicates. The top heatmap demonstrates the difference in fraction of methylation at each gene’s identified differentially methylated region within the promoter, with positive values indicating more methylation in OC104 and negative values indicating more methylation in OC109. Bottom heatmap shows log2 fold change (FC) in gene expression of each gene for OC109 relative to OC104.

### Response of PDOs to carboplatin

PDOs OC104 and OC109 were assessed for response to chemotherapy using drug dose response assay to carboplatin. Despite PDOs being derived from ascites collected within one week of each other from the same patient, responses to carboplatin were distinct (**Figure 6D**). OC104 had an average IC_50_ of 31.92 µM and OC109 had an average IC_50_ of 82.32 µM. Carboplatin has a maximum achievable concentration (C_max_) of 58.4-87.4uM for 15–30Lminutes of carboplatin infusion at AUC 5–6^36–38^. Thus, OC104 is sensitive to carboplatin, whereas OC109 is partially resistant to carboplatin. The divergent *ex vivo* responses are consistent with the heterogenous clinical response noted observed during combination therapy. While a general reduction in the ovarian mass was discernible by imaging, marked tumor burden observed at time of surgery was deemed as unresectable disease. We suspect that OC104 represents the major ovarian mass and that OC109 largely represents the disseminated/chemoresistant disease.

We prioritized several of the genes with differentially methylated promoters for expression analysis to determine whether methylation changes may contribute to resistance phenotypes. Generally, we would expect resistance to be associated with upregulation of oncogenes and down-regulation of TSG, so we considered the directionality of change observed in DMR analysis described above and the general contribution of each gene as an oncogene or TSG based on the COSMIC database as well as literature searches in ovarian cancer. We prioritized tumor suppressor gene HNF1A and oncogenes MDM4, MSI2, and SETBP1, as well as BRCA2 due to its re-expression being associated with resistance in high-grade serous ovarian carcinomas. We isolated RNA from PDOs on two distinct passages and performed qRT-PCR. HNF1A, MSI2, and SETBP1 expression differed in the organoids in directions consistent with change in DMR methylation (**Figure 6E**). MDM4 and BRCA2 did not, suggesting that methylation at these regions does not correlate with expression (**Figure 6E**).

The DMRs at the prioritized genes are shown in **Supplemental Figure 6**.

## Discussion

This study presents a unique case of two PDOs derived from the same patient, with LRS and drug response analyses yielding tumor subtype reclassification. Derivation of two PDOs from a single patient, given otherwise low efficiency rates in generating PDOs from tumor or ascites, suggests the patient-specific nature of PDO generation. While media formulations have been largely developed in endoluminal gastrointestinal cancers, whether they are optimized for success in ovarian cancer PDO development remains uncertain. One recent preprint on mucinous ovarian carcinoma PDOs^15^ modified media formulation compared to that previously developed for ovarian cancer PDOs from Hans Clever’s lab^9^. They found that increased B-27 supplement, Y-27632 ROCK inhibitor and human-EGF, addition of N-2 supplement and FGF-2, and absence of beta-estradiol, heregulin beta-1, and forskolin improved PDO generation. Our formulation was not specifically optimized for ovarian cancer PDOs, with the main difference being the absence of several growth factors and hormones (FGF-10, FGF-2, beta-estradiol, hydrocortisone, gastrin, heregulin-beta1, and forskolin) and addition of MAPK inhibitor SB202190 to inhibit fibroblast growth. Further work will be needed to ensure that PDO conditions are optimal for ovarian cancer, in addition to other tumor features that influence PDO efficiency rate.

Long-read sequencing of cancer genomes remains in its relative infancy, with only 33 ovarian cancers sequenced in the published literature to date^18–21^. Our work is unique in that we have performed LRS on an ovarian carcinoma with features overlapping high-grade serous carcinoma and the enigmatic mucinous carcinoma, and leveraged several strengths of LRS, examining likely pathogenic single nucleotide polymorphisms, structural variants, and differential methylation. In addition, previous LRS of whole genomes in PDOs are limited in parallel information from manipulation of the PDOs. While one report confirmed ecDNA presence with fluorescence in situ hybridization^22^ and another performed karyotyping, ChIP-seq, and Hi-C^23^, each in esophageal PDOs, our study specifically assessed the response of PDOs to carboplatin as a single agent.

Of note, examination of the genomics of these two isogenic PDOs from a single patient led us to retrospectively reconsider whether mucinous carcinoma would have been a better classification for this tumor. In hindsight, the original finding of WT1 negativity could be regarded as an argument against a diagnosis of high-grade serous carcinoma, which is usually positive for WT1, but in the setting of advanced tumors it is not uncommon to observe isolated immunophenotypic results that do not support the final diagnosis and it was therefore readily regarded as noncontributory. The pattern of spread within the peritoneum is certainly in keeping with the patterns seen in the serous tumors in clinical practice. In hindsight, the additional finding of KRAS mutation in the clinical sequencing, and the patient’s poor outcome and rapid disease progression are more compelling for a mucinous ovarian carcinoma, which are rather rare, and characteristically aggressive and chemoresistant in the setting of disseminated disease^39^. Classification of an ovarian carcinoma is relatively straightforward in the majority of the patients when a generous sample is available/on a surgical specimen; however, it is much more challenging in a small biopsy specimen as many high-grade carcinomas have overlapping morphology and share abnormalities in p53. Notably, we found CDKN2A and broader 9p deletions in our PDOs, with a higher magnitude of copy number loss observed in the more resistant OC109, further consistent with mucinous ovarian carcinoma characteristics. It is unlikely that a more specific diagnosis would have led to a change in treatment in current practice for this patient, but beyond the question of whether these findings could have been regarded differently for this patient, they demonstrate the shortcomings of our current classification system and highlight the utility of molecular findings in characterizing tumors beyond the limitations of light microscopy. Indeed, it is unclear to what extent the role of histological subtyping of gynecologic cancers will evolve as treatments become increasingly driven by specific abnormalities. Our SV finding of an alteration in MUC5AC is consistent with literature suggesting MUC5AC may be a biomarker for mucinous ovarian carcinomas^31–34^.

Perhaps most intriguing is that the two PDOs developed from this particular patient display differential responses to carboplatin, despite derivation from samples collected only one week apart. We suspect that clonal variation contributes to this differential response, as otherwise the two PDOs were generally under the same culture conditions. This clonal variation may be at the genetic or epigenetic level and is likely derived from a heterogeneous tumor population. As our PDOs were derived from the cellular fraction of ascites fluid, we cannot determine the anatomical location from which PDOs were derived. We demonstrated a number of prioritized clonal variants that differ between the two PDOs. A SNV in NCOR2, p.Arg1625Cys, found only in OC104 is in a poorly annotated region of the protein, so its effect on protein function is unknown but may interact with the SMC1 domain, influencing how it interacts with other proteins. This heterozygous variant has also been observed in an endometrioid carcinoma of the endometrium (from COSMIC), which of note, is associated with mucinous differentiation and an advanced endometrioid ovarian tumor could also explain the findings in this case, again highlighting the shortcomings of histologic classification. The deletion within BMX is predicted to disrupt much of the N-terminal portion of the protein, which includes the Bruton’s tyrosine kinase pleckstrin homology domain, which has roles in tethering BMX to membranes through interaction with phosphatidylinositol lipids^40^. BMX is typically lowly expressed in epithelial cells but may become activated during cell stress to facilitate wound healing programs^41^. While it is unclear how the chromoplexy involving chromosomes X, 6, and 21 influences tumor biology, it is only observed in OC109, indicating structural rearrangement in clonal variants within this patient. Two of the break points lie within genes, including ZNF280C and HEMK2. HEMK2 encodes N(6)-adenine-specific DNA methyltransferase, which has controversial roles as a DNA or protein methyltransferase^42–45^. ZNF280C is not well studied but may act as a transcription factor. Both variants lie within introns and are predicted to disrupt protein expression. Perhaps most interesting are findings from differentially methylated promoters. Most notably, BRCA2 promoter methylation decreased from 85% methylated in OC104 to 14% methylated in OC109, which is consistent with homologous recombination deficiency being associated with a more favorable response to platinum-based chemotherapy in both our data and in the clinic^46,47^. The patient tested negative for HRD in clinical genomics testing, but this perhaps highlights the importance of repeat samplings in complex presentations of disease. Additionally decreased promoter methylation to the growth factor receptors FGFR2 and ERBB4 occurs in the more platinum-resistant OC109, suggesting growth factor signaling may also influence resistance.

Interestingly, 87% of all differentially methylated promoters were more methylated in the PDO that was more sensitive to carboplatin. We also observe this in high-grade serous ovarian carcinoma cell lines using both isogenic pairs and non-isogenic platinum-sensitive and -resistant lines (data subject of another manuscript in preparation). Promoters are known to be predominantly hypermethylated in cancer relative to their cells of origin^48^, but the literature generally disagrees as to how methylation of promoters changes with chemoresistance. One study using an patient-matched cell line system similarly demonstrated that 84% of differentially methylated loci using reduced-representation bisulfite sequencing are less methylated in the resistant cell line^49^. However, several studies using tumor tissue, which may be heterogeneous in cell makeup, have shown the opposite: chemoresistant tumors gain promoter methylation^50–52^. Further work is required to determine whether the discrepancies are due to heterogeneous nature of sample inputs, or technically driven by method of DNA methylation measure or bioinformatic analysis approach. Definitive investigation is important to fully understand the general failures of DNA methylation inhibitors in ovarian cancer clinical trials^53–64^.

From our gene expression analysis of prioritized genes, we demonstrated that HNF1A, MSI2, and SETBP1 expression correlated with promoter methylation status when comparing the two PDOs in a direction that would be expected to be associated with resistance. HNF1A is interesting given that it is also expressed in gastrointestinal tumors with mucinous features^65^ and HNF1A expression decreases with cancer progression in pancreatic cancer^66^, though its expression increases in cancer stem cells^67^. Little is known about its role in ovarian cancers, though its homolog HNF1B is proposed as a biomarker for ovarian clear cell carcinomas^68^ and may play a role in expression of Folate Receptor alpha^69^. MSI2, or Musashi RNA binding protein 2, is highly expressed in ovarian cancers, has been shown to increase proliferation and growth, and is increased with paclitaxel resistance and in cancer stem cell populations^70,71^, and has been implicated in other female cancers^72^. SETBP1 functions as part of a complex that binds to AT-rich DNA and promotes gene expression^73^ and promotes cancer stemness phenotypes of ovarian cancer cells^74^. Further mechanistic work is needed to demonstrate whether these genes are involved in the carboplatin response differences between our PDOs. Interestingly, we expected the loss of methylation of BRCA2 in the more platinum-resistant OC109 to be associated with treatment response, as loss of homologous recombination is known to confer platinum sensitivity^75–77^. However, we did not observe that the loss of methylation at the loci identified in our analysis is associated with expression changes, indicating that this part of the promoter is non-essential for BRCA2 regulation.

While the depth of investigation using LRS is a strength of the study, the fact that only one pair of PDOs derived from a single patient was investigated is also a weakness as to the generalization of these research findings. It would be valuable to investigate the findings here in a broader set of mucinous ovarian carcinoma PDOs, which have been recently reported^15^. Additionally, we can only speculate as to the involvement of specific SNVs, SVs, and DMRs in the biology of these tumors. Manipulating the genetics of PDOs remains a challenge in the field and requires much more extensive studies. Future work will be needed to assess how these findings impact ovarian carcinoma biology and treatment response.

## Acknowledgements

This work was supported by the Department of Defense Ovarian Cancer Research Program (OC240130) and a Wisconsin Partnership Program New Investigator Award to J.D.L. We thank Mark Burkard’s laboratory for providing CAL51 cells and for valuable advice on DNA content analysis. We acknowledge the Center for Precision Medicine for access to the PromethION platform and the University of Wisconsin Biotechnology Center for support with the long-read sequencing pipeline. The author(s) thank the University of Wisconsin Carbone Cancer Center BioBank, supported by P30 CA014520, for use of its facilities and services. The authors wish to thank the University of Wisconsin Translational Research Initiatives in Pathology (TRIP) Laboratory, supported by the UW Department of Pathology and Laboratory Medicine, UWCCC (P30 CA014520) and the Office of The Director- NIH (S10 OD023526) for PDO fixing, embedding, and staining services.

## Data Availability Statement

Long-read sequencing data will be deposited to dbGap (phs004795.v1.p1). Processed pipeline outputs and code is available upon request.

---

**Supplemental Figure 1.**
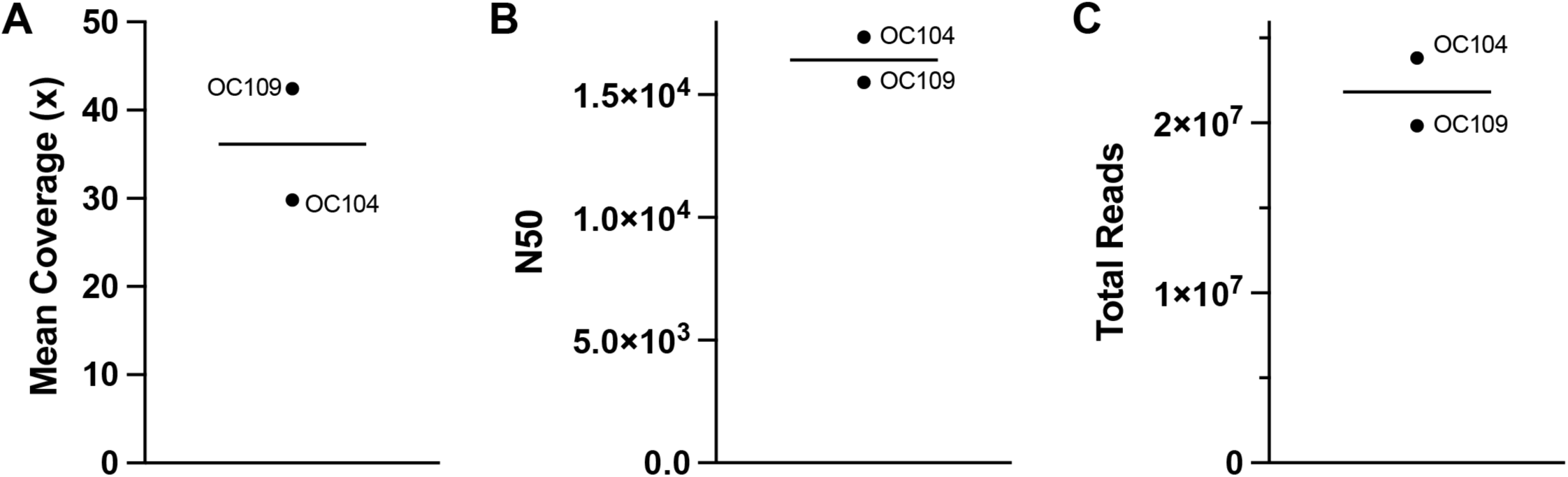
Long-read sequencing metrics. (A) Mean coverage across the genome. (B) N50, or the read size for which the sum of it and all larger reads make up half of the total number of base pairs sequenced. (C) Total number of reads for each sample. For each figure panel, individual data is shown for each sample (PDO name labelled) and the mean is represented by the horizontal bar.

**Supplemental Figure 2.**
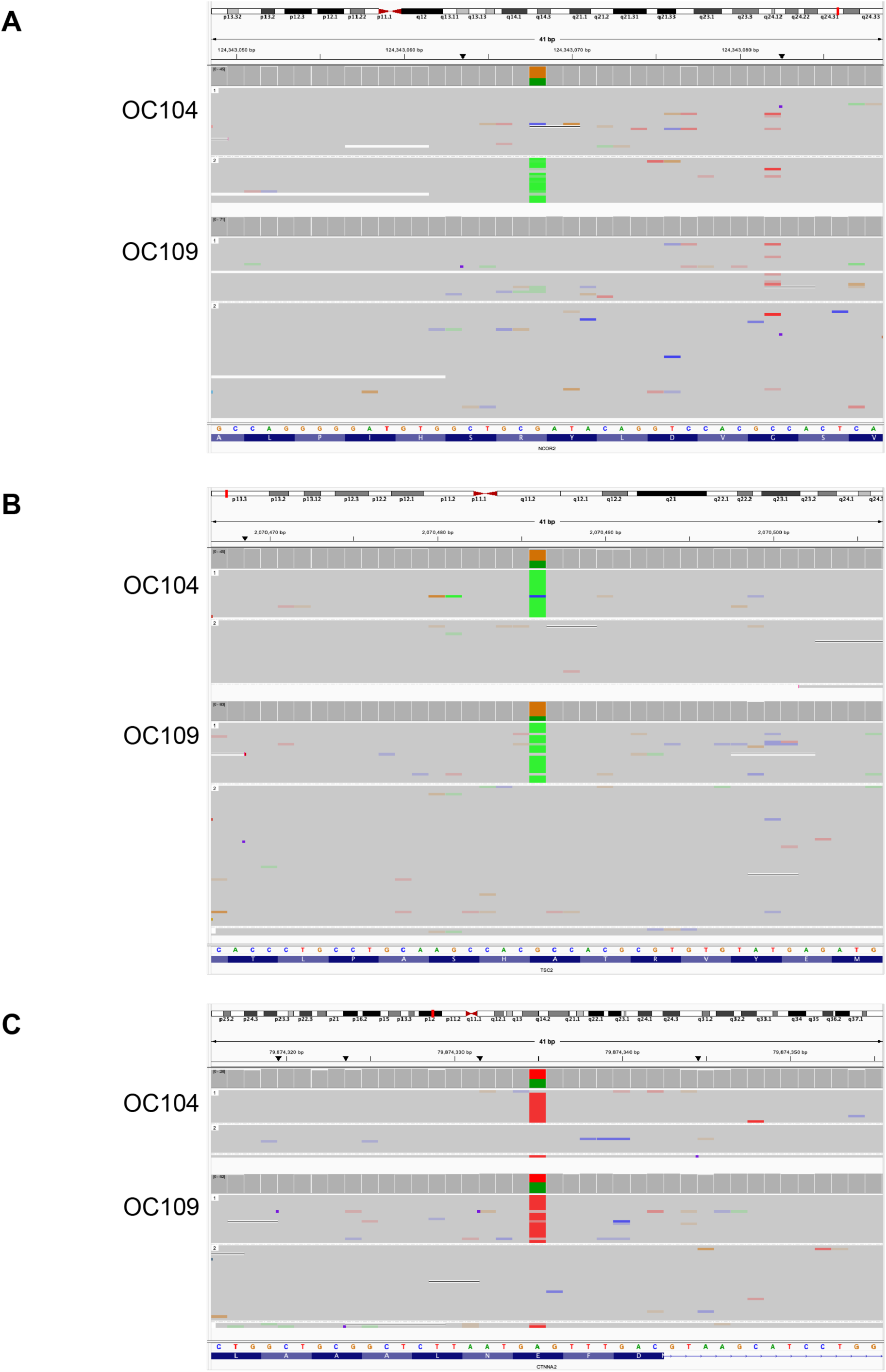
IGV diagrams of LRS single-nucleotide polymorphism nonsynonymous variant calls from PDOs. (A) NCOR2 p.Arg1625Cys caused by a G>A nucleotide variant. (B) TSC2 p.Ala583Thr caused by a G>A nucleotide variant. (C) CTNNA2 p.Glu282Val caused by an A>T nucleotide variant. For each panel, reads are phased into haplotypes 1 and 2 or unassigned. Haplotypes are randomly assigned.

**Supplemental Figure 3.**
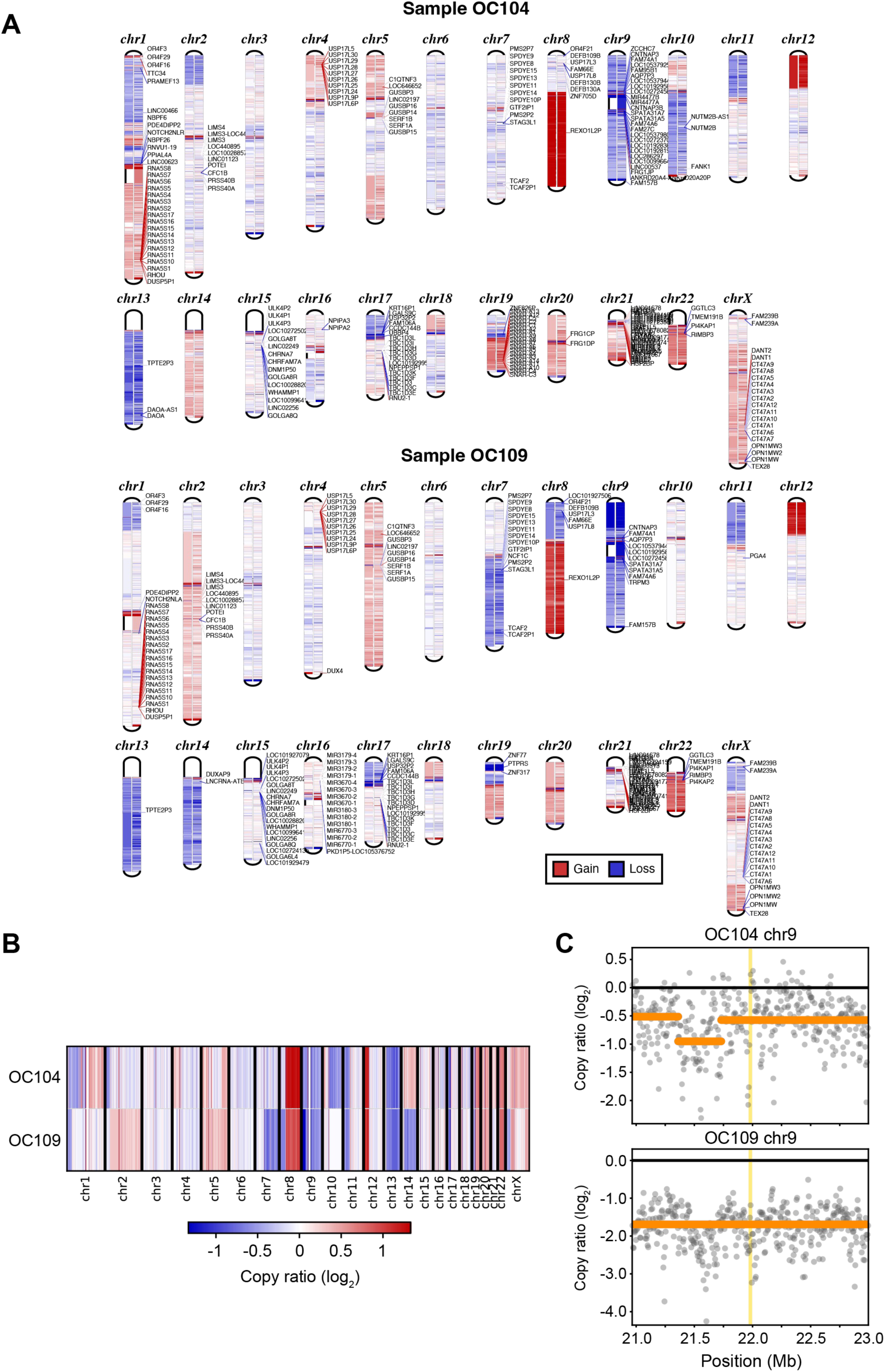
Copy number analysis of PDOs. (A) Copy number across genome of PDOs, Top: OC104, Bottom, OC109. For each chromosome, segment copy number ratios are shown on the left side, and copy number ratio by bin on the right side. Genes with an absolute log2 fold change >2 are labelled. (B) Side-by-side copy number ratios of PDOs. (C) Chromosome 9p locus that contains the CDKN2A gene (yellow vertical bar). Gray points represent copy number bin values, and orange horizontal lines indicate the segment level copy number ratio.

**Supplemental Figure 4.**
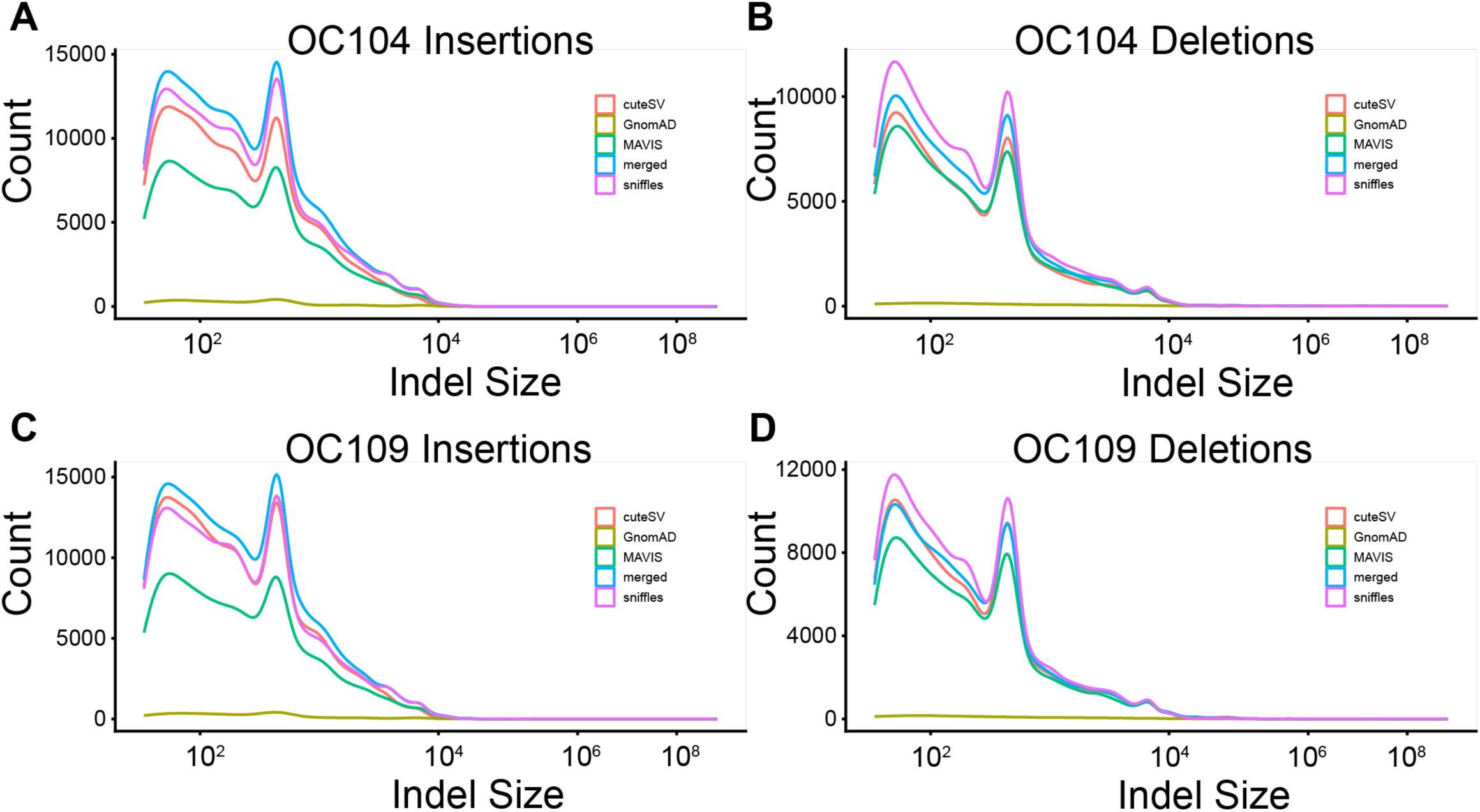
Distribution of lengths of insertions and deletions (Indels) by step in the filtering process. (A) OC104 insertions, (B) OC104 deletions, (C) OC109 insertions, and (D) OC109 deletions. Indel length is described in base pairs. CuteSV indicates indels called by the cute SV SV caller. Sniffles indicates indels called by the sniffles2 SV caller. Merged indicates the SVs merged by MAVIS, prior to filtering. MAVIS indicates the SVs after MAVIS’s filtering step. GnomAD indicates those SVs that passed filters removing regions of very high or very low coverage in the population.

**Supplemental Figure 5.**
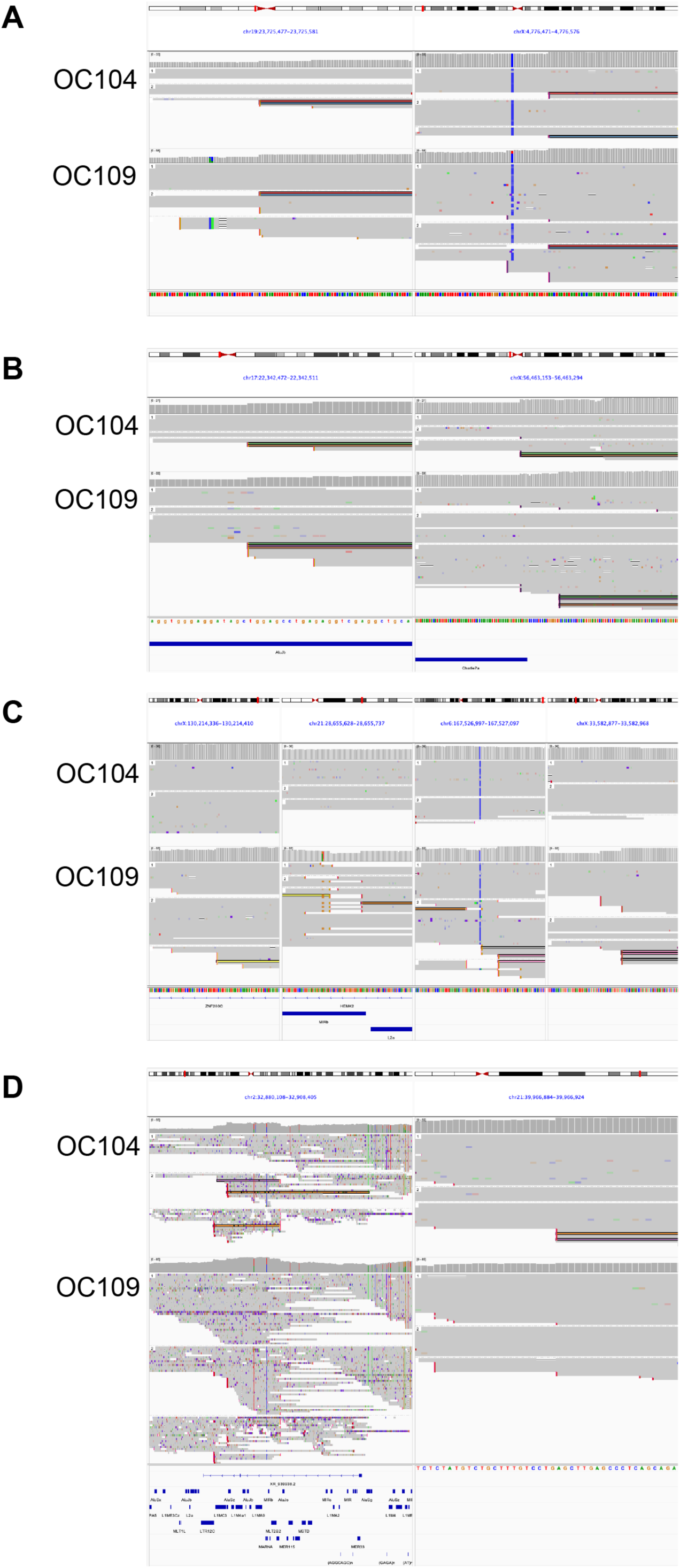
IGV diagrams of LRS translocation calls from PDOs. (A) Breakpoint locations for BND1 between chr19 (left) and chrX (right). (B) Breakpoint locations for BND2 between chr17 (left) and chrX (right). (C) Breakpoint locations for BND3 between chr X (far left), chr21 (middle left), chr6 (middle right), and chrX (far right). (D) Breakpoint locations for BND4 between chr2 (left) and chr21 (right). Annotated genes and RepeatMasker repeat regions are shown below alignment tracks. For each panel, reads are phased into haplotypes 1 and 2 or unassigned.

**Supplemental Figure 6.**
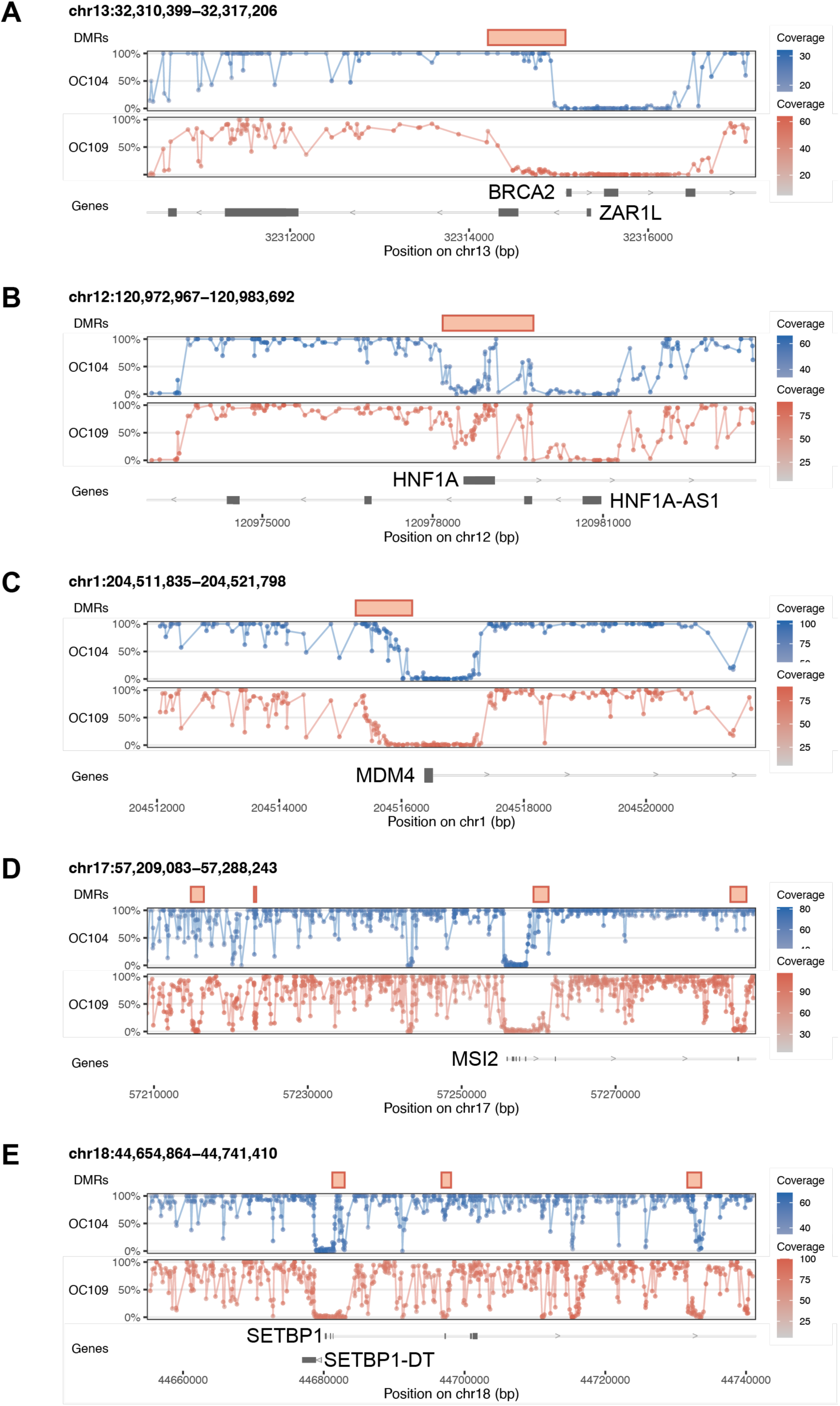
Signal tracks of differentially methylated promoter regions of (A) BRCA2, (B) HNF1A, (C) MDM4, (D) MSI2, and (E) SETBP1 showing 5-methyl cytosine methylation, the detected DMR, and the gene annotation track.

**Supplemental Table 1**. Organoid media formulations, suppliers, and catalog numbers.

**Supplemental Table 2**. Single nucleotide variants prioritized from long-read sequencing of OC104 and OC109 PDOs with database annotations. Each SNV is annotated with effect predictions (AlphaMissense, PolyPhen, SIFT) with the category with score in parentheses. The total number of ovarian cancer patients in the AACR GENIE database with the exact variant are shown, as well as breakdown by subtype for either exact variant match or any alteration to the gene. Genes are also annotated for presence/frequency in COSMIC (occurrence in cancer) and GnomAD (frequency in population) databases. AF = allele fraction, TSG = tumor suppressor gene, OG = oncogene.

